# Cooperative DNA binding by a splice variant of the transcription factor ETS1 that lacks autoinhibition

**DOI:** 10.1101/070185

**Authors:** Daniel Samorodnitsky, Courtney Szyjka, Gerald B. Koudelka

## Abstract

ETS1 is the archetypal member of the metazoan ETS transcription factor family. All the members of this family bind to the conserved GGA(A/T) core ETS binding sequence (EBS). Several ETS family members exhibit autoinhibition, wherein one domain of ETS blocks its own DNA-binding activity until certain conditions are met. Relief of autoinhibition in these proteins is coupled to homo- or hetero-dimer formation. Relief of autoinhibition in full-length ETS1 is catalyzed by non-specific interaction with DNA, which facilitates the formation of protein dimers.The ETS1 splice variant, ETS1-p42, lacks an exon that encodes a crucial part of the autoinhibitory module. Thus,ETS1-p42 does not autoinhibit. We showed that the absence of autoinhibition allows ETS1Δ335, an N-terminal deletion that recapitulates the DNA-binding of ETS1-p42, to form homodimers in the absence of DNA. However, ETS1-p42 is thought to bind its DNA sites only as monomers. Here, we explore this paradox and show that ETS1Δ335 binds cooperatively to DNA containing two EBSs. We also show that residues in ETS1’s DNA recognition helix mediate this DNA-binding cooperativity. Furthermore we also find that a single EBS can bind two ETS1Δ335 subunits. Finally, we show that DNA acts catalytically to separate unbound ETS1Δ335 dimers into monomers. Together,these results suggest a model of ETS1-p42 DNA-binding, where ETS1-p42 can bind to a single EBS with 2:1 stoichiometry.

## Introduction

Precise regulation of gene expression is required for normal development and function of all biological entities. Gene expression is commonly regulated through the action of sequence-specific DNA-binding transcription factors. Multiple transcription factor binding sites frequently occur in a single regulatory element, allowing interaction and potential modulation of the behavior of these factors. The cooperative binding of multiple factors within a single promoter allows for the generation of highly complex, precisely controlled gene regulatory circuits. The ability of a given protein to interact with other gene regulatory proteins and form a variety of transcription factor complexes is determined by the number and type of structural domains possessed by each protein.

In addition to domains that permit protein-DNA interaction and protein-protein docking, autoregulatory domains may downregulate transcription factor complex activity by blocking or altering DNA binding activity and/or inhibiting oligomerization. In order for the transcription factor to become fully active, these domains may be required to change their structure, be removed via alternative splicing and/or be post-translationally modified (1).We are interested in exploring the mechanisms by which autoregulatory domains modulate transcription factor function. Since there is an exact correspondence between its DNA binding affinity and transcriptional regulatory activity, for these studies we are investigating the effect of the autoinhibitory domains of ETS1 on its oligomerization and DNA-binding properties.

ETS1 is the archetypal member of the ETS transcription factor family (2). The name “ETS” is an acronym for “E-twenty-six specific”, for the avian retrovirus whose genome contains an oncogenic variant of a cellular ETS protein (3). The ETS family is conserved throughout the metazoan lineage (4-6). ETS proteins specifically bind DNA via the ETS domain, a variant of the “winged” helix-turn-helix motif (7-9). The domain itself is a conserved ~85 amino acid residue region, consisting of a series of three α-helices, H1, H2, and H3, appended to a four-strand β-sheet. Because all ETS proteins share this DNA binding motif (10), each binds the ETS binding sequence (EBS), GGA(A/T). The full-length isoform of ETS1 contains several domains that regulate both its DNA-binding affinity and its ability to form a variety of multimeric DNA-binding complexes (11–16). In the autoinhibited form, two of these domains, the N-terminal (NTD) and C-terminal (CTD) autoinhibitory domains flank the ETS domain and fold over helix H1 of the ETS domain (12,17). Notably, helix H3, which is responsible for directly contacting the EBS, is not blocked and is solvent accessible in the autoinhibited form of ETS1.

Previously, the autoinhibitory module was thought to provide a “graded control” of ETS1 DNA-binding affinity, allowing ETS1 the ability to produce different levels of transcriptional regulation depending on the bound DNA sequence and protein partner(s) present (15,18). However, our recent work showed that autoinhibition also functions to block ETS1 dimerization in the absence of DNA, suggesting that autoinhibition functions to “prime” ETS1 for non-specific interaction with DNA, and, in turn, interaction with a partner protein (2). When the ability to autoinhibit is lost, either as a result of mutation or the splicing out of exon VII, ETS1 dimerizes spontaneously (2).

A major splice variant of ETS1, termed ETS1-p42, lacks exon VII that encodes the N-terminal inhibitory domain. As a result, ETS1-p42 is not autoinhibited. This indicates that the NTD is crucial for ETS1 autoinhibition. ETS1-p42 reportedly binds to a head-to-head site containing two EBSs in the promoter for the matrix metalloproteinase MMP3 (12,14). However, despite the presence of two EBSs, ETS1-p42 apparently binds as a monomer to the MMP3 promoter, whereas the full-length ETS1-p51 variant seemingly binds this sequence only as a homodimer. (12). Thus, in addition to autoinhibition, exon VII also apparently plays a role in modulating ETS1 homodimerization (2). Furthermore, despite binding as an apparent monomer, ETS1-p42 needs both EBSs to drive expression from an MMP3-luciferase reporter construct (14,19). To date these contradictions have not been explored.

Taken together, these findings raise a number of important questions. Does ETS1-p42 bind DNA as a monomer? If so, why are multiple EBSs needed for ETS1-p42 transcriptional activation? Furthermore, if a consequence of losing autoinhibition is ETS1-p42 self-association in the absence of DNA, what regions of the protein are responsible for mediating this self-association? In this study, we find that despite its inability to autoinhibit, ETS1-p42 binds cooperatively to a promoter sequence containing two EBSs. Additionally, footprinting results indicate that two ETS1-p42 proteins bind the MMP3 promoter, potentially explaining the necessity for both EBSs to drive expression. Finally, we show that in the absence of an intact autoinhibitory module, ETS1-p42 appears to interact with DNA in a fashion where DNA induces a rearrangement of a free ETS1-p42 dimer into a monomer that is competent to bind DNA. This behavior differs markedly from that of ETS1-p51, where DNA induces a monomers to dimers transition (2).

## Methods

### Proteins

Protein purification was performed as described in (2). Sequences of DNA primers used for R394A/Y395A and R378C mutagenesis can be found in (2) as well. The sequences of the primers for L337A mutagenesis are:

> F: 5’ CCCCATGGCTATCCAG<u>GCA</u>TGGCAGTTTCTTCTGG 3’
>
> R: 5’ CCAGAAGAAACTGCCA<u>TGC</u>CTGGATAGCCATGGGG 3’

The altered bases are underlined.

### DNA Purification

Sequences containing naturally occurring EBSs were ordered from Integrated DNA Technologies (Coralville, IA) and purified as described in (2).

### EMSA

Double-stranded DNA was radiolabeled by incubating DNA with γ - [^32^P]ATP (6000 Ci/mmol) (PerkinElmer Life Sciences) in the presence of T4 polynucleotide kinase (Epicenter, Inc., Madison, WI). EMSA was performed and analyzed as described in (2).

### DNase I Footprinting

The preparation of DNA templates, the footprinting assays, and resolution of cleavage products was performed as described in (2).

### Protein Crosslinking

Crosslinking assays were performed as described in (2).

### FRET Analysis

5mg of purified ETS1Δ335 stored in 1X SPB_50_ (3.8mM NaH_2_PO_4_ plus 16.2 mM Na_2_H_2_ PO_4_), pH 6.8 and 500mM NaCl was labeled by incubating with 1 mg of either Alexa Fluor 488 succinimidyl ester (A-2000, Thermo Scientific) or dabcyl succinimidyl ester (D-2245, Thermo Scientific) for three hours at 4°C in the dark with rocking. At pH 6.8-7.0, amine-reactive labeling is biased towards the N-terminal amine. The reaction was quenched by successively dialyzing the sample against fresh 1X SPB_50_ and 500mM NaCl. Successful labeling was quantified according to manufacturer’s instructions. The activity of the labeled protein was assessed by examining its ability to bind MMP3 DNA in an EMSA (data not shown).

The association states of the labeled proteins were assayed using an LS 50B Luminescence Spectrometer (PerkinElmer Life Sciences). 130 μMol of Alexa Fluor 488-labeled ETS1Δ335 was incubated in 100uL of 1X SPB_50_ and 500 mM NaCl at 4°C for 20 minutes. This reaction was excited at 490nm, using a 4nm slit width. The emission was measured at 525 nm using a 6nm slit width. Measurements were taken every second for 10 seconds. Following this, the fluorescence of a mixture of 130 μMol Alexa Fluor labeled and 130 μMol dabcyl labeled ETS1Δ335 was measured. To this reaction, increasing amounts of DNA were added until saturation was reached. After each titration, the reaction was mixed and allowed to incubate at 4°C for 20 minutes. This assay was repeated five times for each of the three tested DNA sequences. Results presented are averages of all points taken across all five repetitions.

## Results

### ETS1-p42 binds cooperatively to the MMP3 promoter

To begin examining the role of autoinhibition in governing the DNA binding mode of ETS1-p42, we determined the DNA binding affinities of ETS1Δ335. ETS1Δ335 contains all residues necessary for DNA-binding, as well as the C-terminal inhibitory domain, and thus recapitulates the DNA-binding activity of ETS1-p42 (14,19). Importantly, and identical to ETS-p42, the ETS1Δ335 protein lacks the N-terminal inhibitory domain encoded by exon VII. ETS1Δ335 also bears a deletion of the N-terminal PNT and transactivation domains present in ETS1-p42. However, these domains do not contribute to DNA binding, dimerization, or autoinhibition (14,19). All experiments described here use ETS1Δ335 as a surrogate for ETS1-p42. We began by determining the affinity of ETS1Δ335 for the EBSs in the MMP3 promoter, which controls expression of the matrix metalloproteinase, stromelysin-1, and in the promoter for p53, the DNA damage-sensing protein. Both promoters contain two EBSs arranged in a head-to-head fashion, separated by four bases. The sequences of the EBSs contained in each promoter are identical, but the flanking sequences outside of the EBSs differ (see Table 1 for sequences). Additionally, we determined the affinity of ETS1Δ335 for MMP3 M1 and MMP3 M2, which are identical to MMP3 except that each bears a mutation in one or the other of the two EBS that prevents stable dimerization of the full length ETS1 isoform (14).

The affinity of ETS1Δ335for each sequence was determined by electrophoretic mobility shift assay (EMSA). We find that when increasing concentrations of ETS1Δ335 are added to radiolabeled DNA containing either the MMP3 EBS, ETS1Δ335 forms a single complex with this DNA (Figure 1). As measured by EMSA, the K_D_ of the ETSΔ335-MMP3 complex is 1.2 nM (Table 1). The ETS1Δ335 – MMP3 complex is smaller than that formed by the full-length ETS1Δ280 variant (2). ETSΔ280 binds to the MMP3 site as a dimer. This finding suggests that, despite containing two EBSs, a single ETS1Δ335 monomer binds to MMP3 in EMSA. Although our results support previous work showing that ETS1Δ335 binds to MMP3 as a monomer (12), we find that mutating one or the other of the two EBSs reduces the affinity of ETS1Δ335 for the promoter sequence (Table 1). Regardless of the number of binding sites present, the same sized complex was formed.

**Figure 1:**
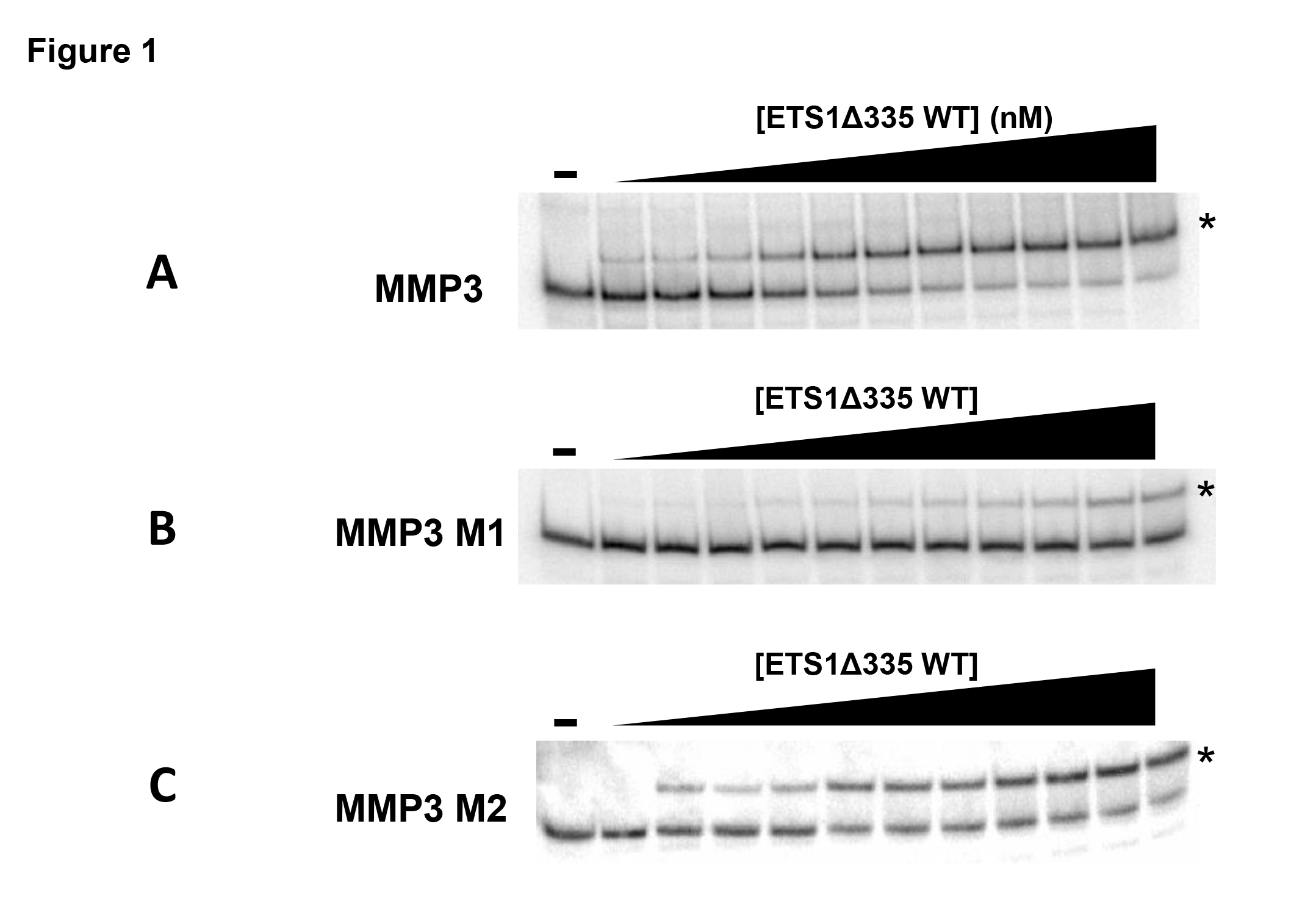
ETS1Δ335-DNA complex formation. Increasing concentrations of ETS1Δ335 were mixed with radiolabeled binding site containing, either MMP3 WT (A), MMP3 M1 (B), or MMP3 M2 (C), (see Table 1for sequences) and complexes resolved via native PAGE as described in Methods and Materials. Shown is a Phosphorimager scan of the gel. Protein-DNA complexes are denoted with asterisks. “-”indicates no protein added. The concentration in the first titration lane for all three gels is 0.0285 nM, and is increased in step-wise fashion. The final lane contains a concentration of 28.5 nM.

**Table 1:**
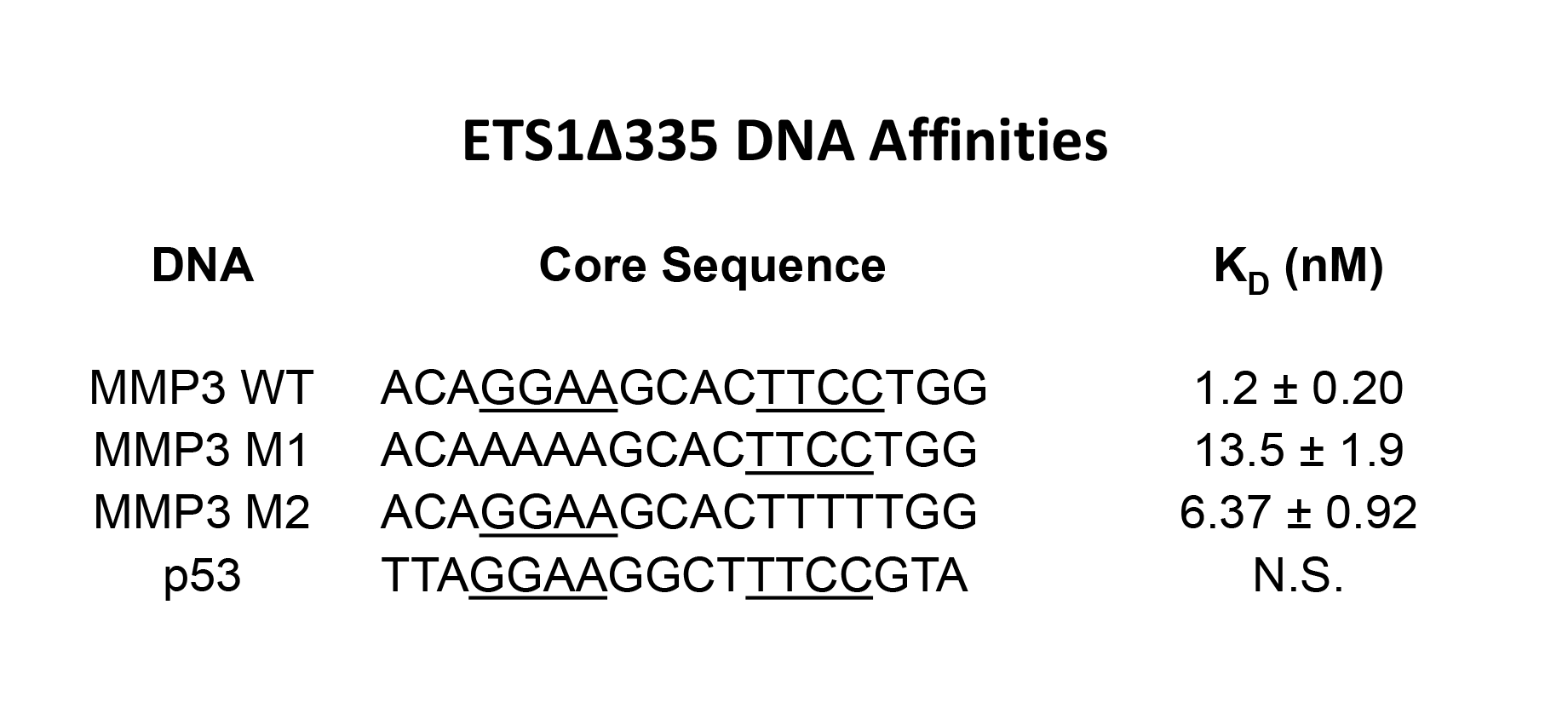
Affinity of ETS1A335 for wild-type and mutant ETS1 binding sites. The sequences and affinity (dissociation constants, K_D_) +/- standard deviation of ETS1A335 for naturally occurring and mutantbinding sites were determined by EMSA (see Methods and Materials). EBSs present in each sequence are underlined.“N.S” is “non-specific”, where ETS1A335 formed no identifiable complex with the DNA site.

Although p53 promoter sequence contains two EBSs whose sequences are identical to that of MMP3, it does not form an EMSA-detectable protein-DNA complex with ETS1Δ335. The inability of ETS1Δ335 to bind to the p53 site is further evidence of the importance of indirect readout in determining the affinity of ETS proteins for its binding sites (2).

Despite seemingly binding as a monomer, the loss of affinity observed upon mutating one or the other EBS in the MMP3 promoter sequence suggests that ETS1Δ335 cooperatively binds this DNA. To help resolve this paradox we probed ETS1Δ335 -DNA complex formation by DNase I footprinting. Added ETS1Δ335 completely protects both EBS core sequences in wild type MMP3 from digestion by DNAse I (Figure 2A). Careful inspection of Figure 2A reveals that the ETS1Δ335 -mediated protection of the 5’ EBS appears at lower protein concentration than the 3’ EBS. Thus, although our EMSA results suggest that ETS1Δ335 binds the MMP3 promoter as a monomer, these findings indicate that apparently this protein binds this DNA as a cooperative dimer. Consistent with the suggestion that ETS1Δ335 binds DNA cooperatively, we find that ETS1Δ335 protects both EBSs in the MMP3 M1 and M2 mutant sequences, despite the fact that each of these sites bears multiple mutations in one of the two EBS present in the wild-type MMP3 DNA site. (Figure 2B & Figure 2C).

**Figure 2:**
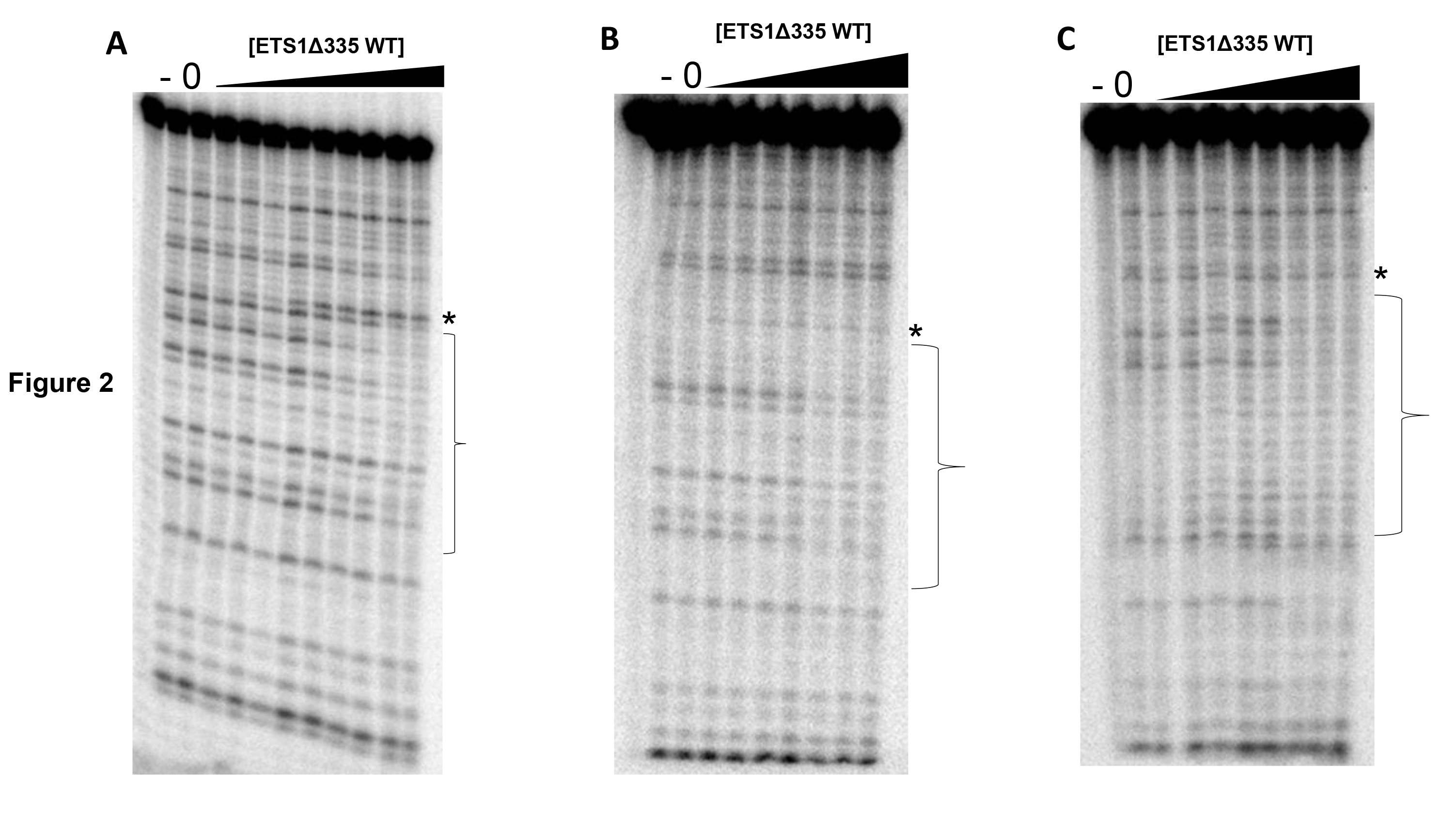
DNase I footprints of ETS1A335-DNA complexes. Radiolabeled DNA bearing MMP3 wild type (A), MMP3 M1 (B), or MMP3 M2 (C) were incubated with increasing amounts of ETS1Δ335. Complexes were then subjected to DNase I cleavage and the resulting DNA fragments were resolved via denaturing PAGE. Shown is a Phosphorimager scan of the gel. In all three panels the lane labeled with a ‘-’ contains uncleaved DNA and the lane labeled with 0 contains DNA incubated with DNAse I in the absence of protein. Positions of bases protected from cleavage by ETS1Δ335 binding are bracketed. Enhancements of cleavage are noted with asterisks.

### ETS1Δ335 Dimerizes in the Absence and Presence of DNA

Having established that ETS1Δ335 cooperatively binds EBS-containing DNA, we wished to explore the oligomeric state of the protein in the absence or presence of various sequence DNAs. To do this we used our previously published sulfhydryl-crosslinking assay (2). In this assay, an R378C mutation is made in ETS1, thereby adding a reactive sulfhydryl that can link individual ETS1 monomers in the presence of a crosslinking reagent. This mutation does not affect the DNA binding specificity of ETS1Δ335. Consistent with our earlier report (2), ETS1Δ335 R378C dimerizes in the absence of DNA (Figure 3). Adding MMP3 WT and MMP3 M1 does not change the amount of crosslinked ETS1Δ335 R378C dimers (Figure 3). Adding MMP3 + 4, a variant that separates the two EBSs present in MMP3 by an extra four bases (2), and to which ETS1Δ335 binds as a mix of monomers and dimers (data not shown) (14), also does not change the amount of dimer formed when compared to no DNA. Similarly, adding either poly dI-dC or the promoter from human p53, which contains two EBSs but to which ETS1Δ335 does not bind (Table 1) also do not affect the amount of crosslinked ETS1Δ335 R378C dimers. However, adding of MMP3 M2 reduces the amount of ETS1Δ335 R378C dimers in comparison to either no DNA or the addition of any other DNA sequence (Figure 3).

**Figure 3:**
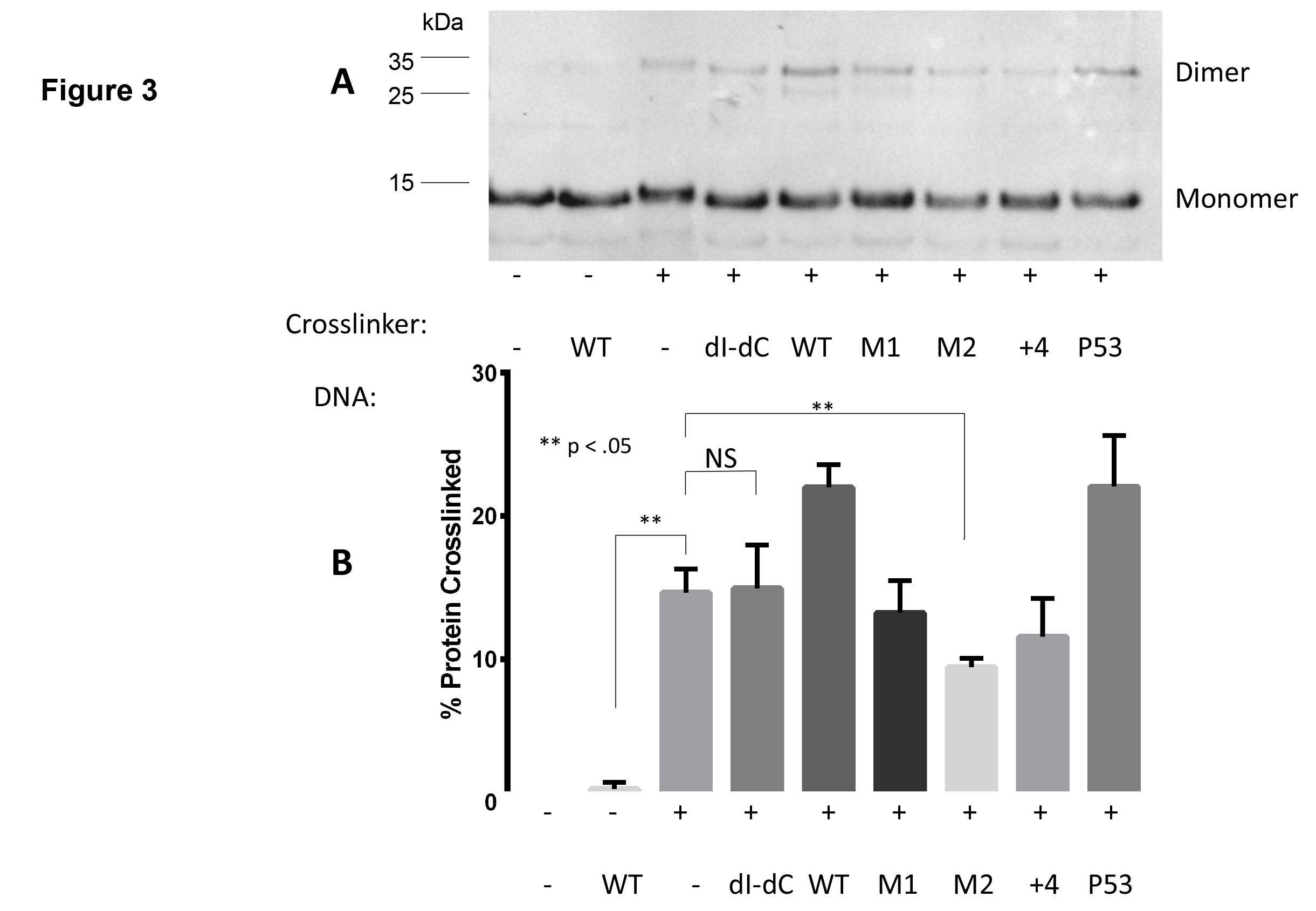
ETS1Δ335 dimer formation. (A) Shown is an immunoblot of ETS1Δ335 R378C crosslinked by BMOE in the absence or presence of the indicated DNA sequence. dI-dC refers to poly dI-dC, a repeating inosine-cytosine oligonucleotide used as a non-specific binding template. The positions of protein monomers and dimers are indicated. Intermediate bands between monomers and dimers are thought to result from incomplete synthesis of ETS1 protein when purified from *E. coli.* (B) Quantification of results seen in (A). Error bars represent standard deviation derived from four or more replicate experiments.

### Cooperative DNA binding by ETS1Δ335 is mediated by helix H3

To further investigate the mechanism of ETS1Δ335 cooperative DNA binding, we determined the DNA binding properties of ETS1Δ335 R394A/Y395A. These residues make two of the three base-specifying contacts between ETS1 and DNA bases within the EBS (12). In ETS1-p51, in addition to blocking DNA binding to the MMP3 promoter, these two mutations disrupt autoinhibition, thereby allowing ETS1-p51 to dimerize in the absence of DNA (2).

Interestingly, despite lacking two DNA contacting residues, ETS1Δ335, R394A/Y395A binds to wild-type MMP3 and the M1 and M2 mutant sites. The affinities of ETS1Δ335 R394A/Y395A for MMP3 wild type and MMP3 M1 are ~10-fold lower than that of wild-type ETS1Δ335 for these sites. The affinity of ETS1Δ335 R394A/Y39A for MMP3 M2 is unchanged from that of wild-type ETS1Δ335 (Table 2). The complexes formed between ETS1Δ335 R394A/Y395A and MMP3 WT, M1, and M2 have identical electrophoretic mobilities. These complexes have similar mobility to ETS1Δ335 WT-DNA complexes, suggesting that the mutations do not alter the stoichiometry of ETS1Δ335-DNA complexes as detected by EMSA (Figure 4A).

**Figure 4:**
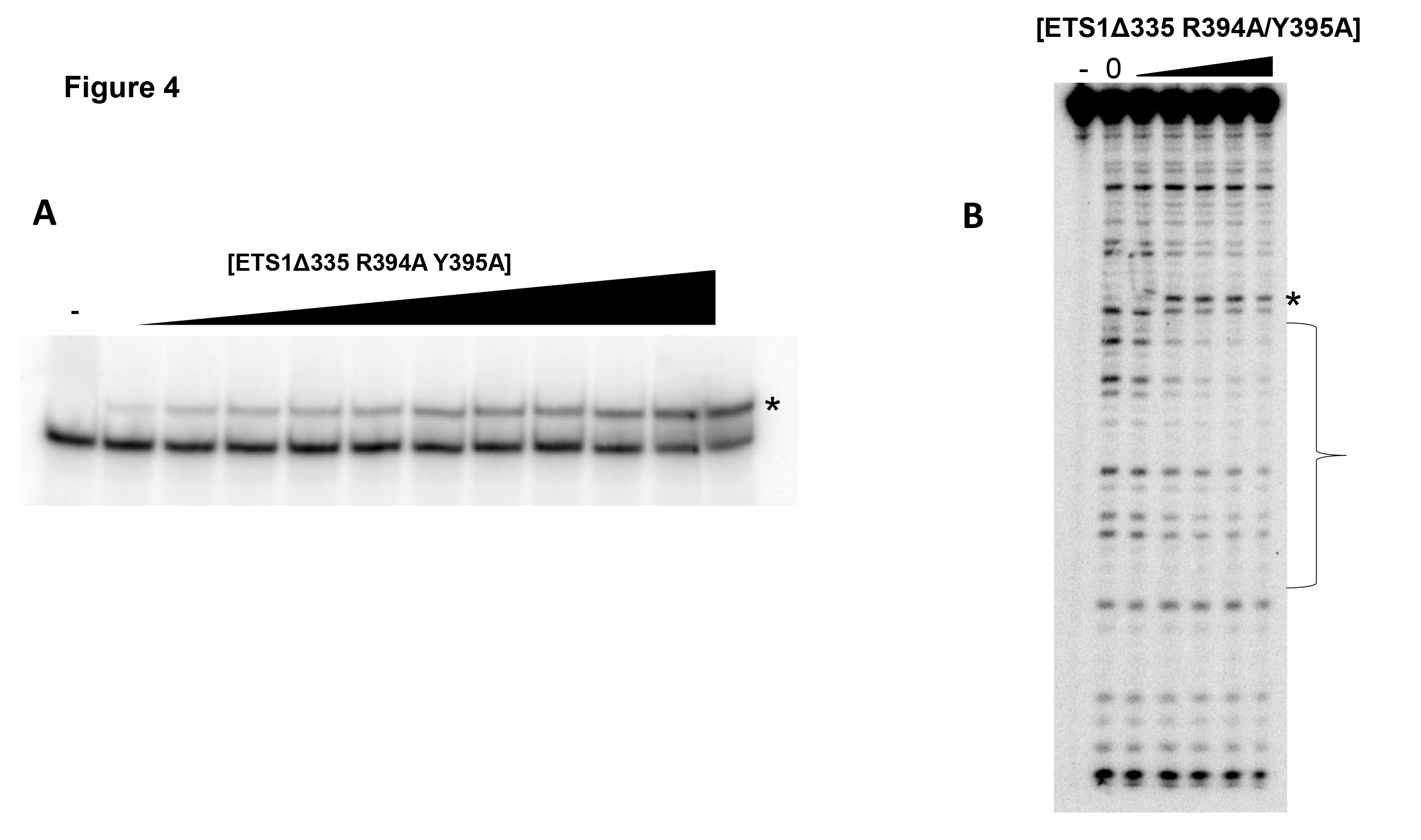
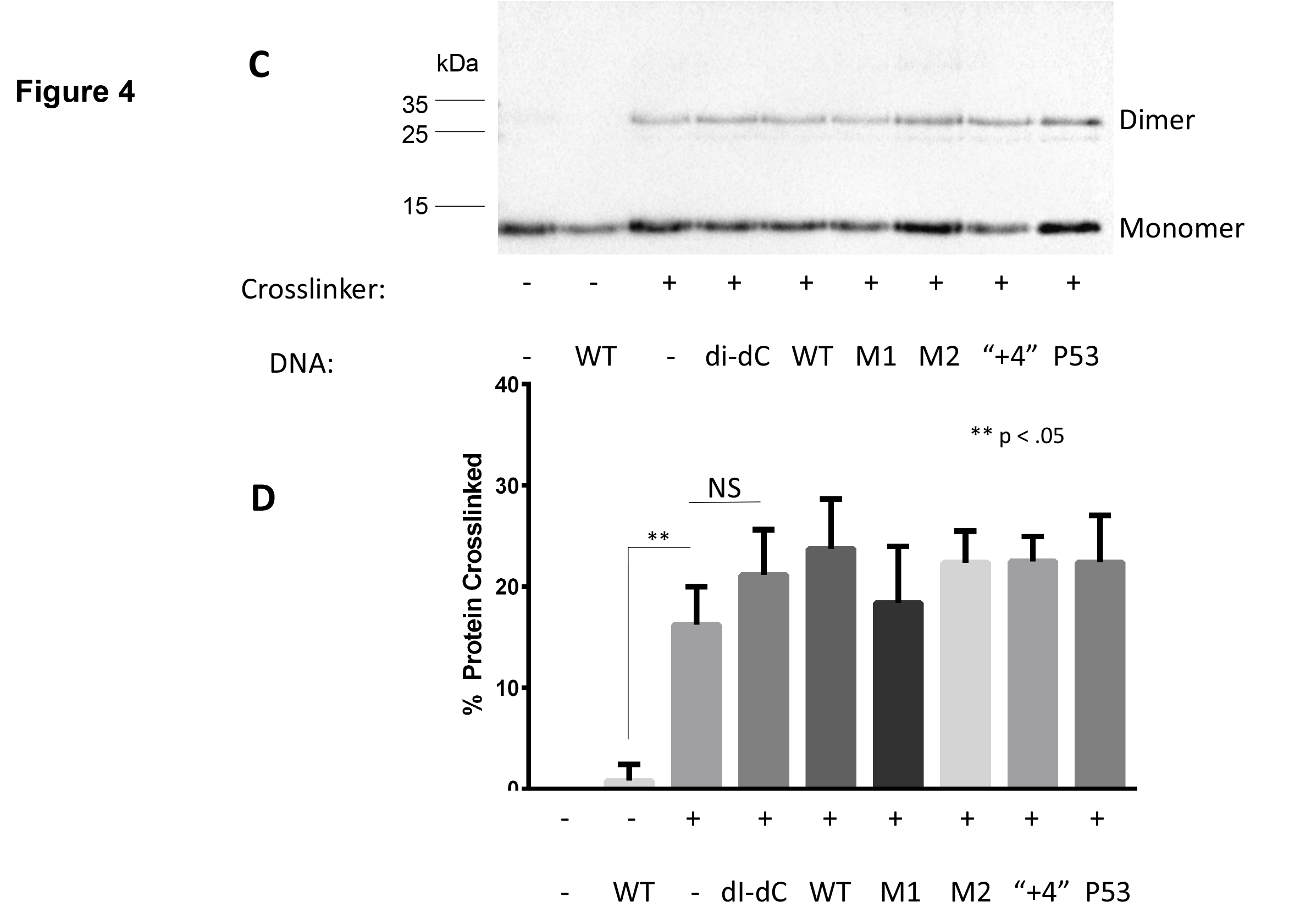
DNA Binding and dimer formation of ETS1Δ335 R394A/Y395A. (A) Increasing concentrations of ETS1Δ335 R394A/Y395A were mixed with radiolabeled binding site containing, MMP3 WT. Protein-DNA complexes were resolved via native PAGE as described in Methods and Materials. Shown is a Phosphorimager scan of the gel. Protein-DNA complexes are denoted with asterisks. “-”indicates no protein added. (B) Radiolabeled DNA bearing MMP3 wild type were incubated with increasing amounts of ETS1Δ335 R394A/Y395A. Complexes were then subjected to DNase I cleavage and the resulting DNA fragments were resolved via denaturing PAGE. Shown is a Phosphorimager scan of the gel. The lane labeled with a ‘-’ contains uncleaved DNA and the lane labeled with 0 contains DNA incubated with DNAse I in the absence of protein. Positions of bases protected from cleavage by ETS1Δ335 R394A/Y395A binding are bracketed. Enhancements of cleavage are noted with asterisks. (C) Shown is an immunoblot of ETS1Δ335 R378C/R394A/Y395A crosslinked by BMOE in the presence or absence of the indicated DNA sequences. Positions of protein monomers and dimers are indicated. (D) Quantification of (C). Error bars represent standard deviation derived from four or more replicate experiments.

**Table 2:**
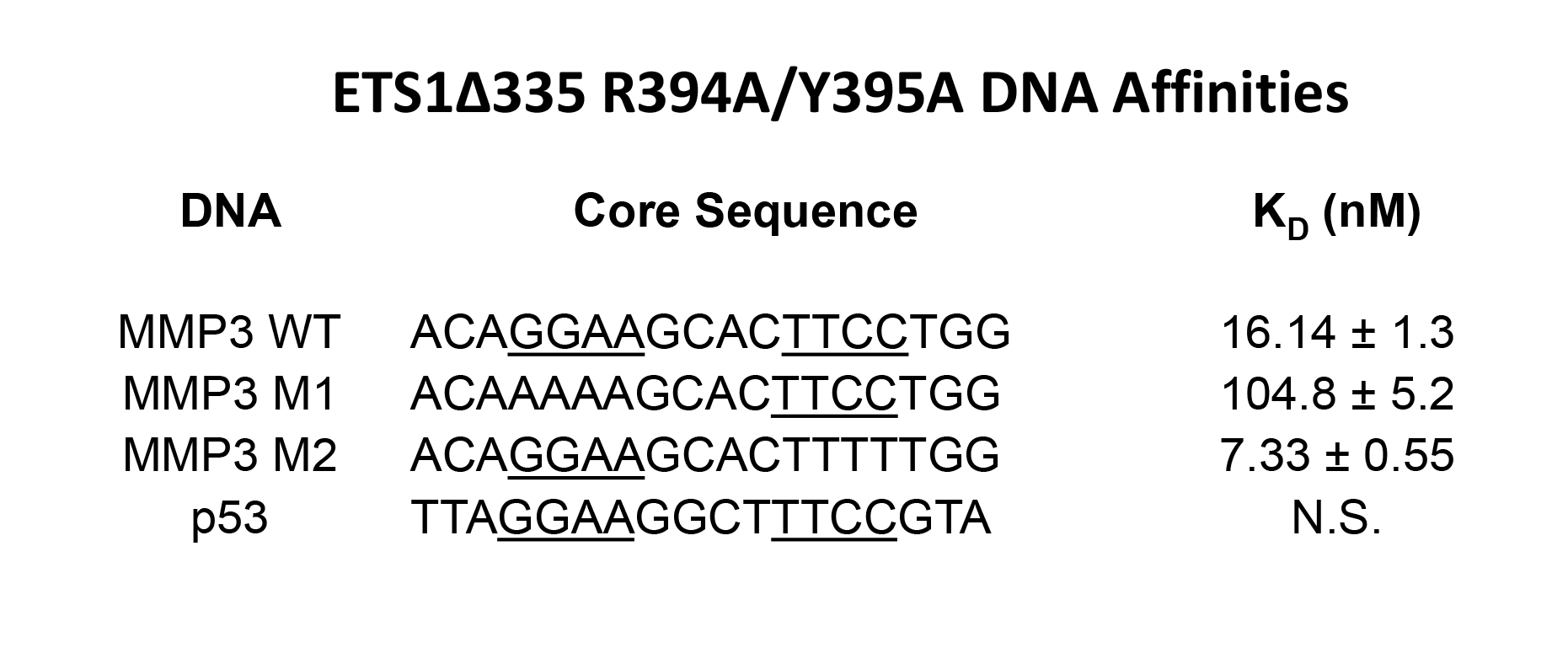
Affinity of ETS1Δ335 R394A/Y395A for different DNA sequences. The sequences and affinity (dissociation constants, K_D_) +/- standard deviation of ETS1Δ335 R394A/Y395A for naturally occurring and mutant binding sites were determined by EMSA (see Methods and Materials). EBSs present in each sequence are underlined. “N.S” is “non-specific”, where of ETS1Δ335 R394A/Y395A formed no identifiable complex with the DNA site.

We used DNAse I footprinting to further examine the effect of the R394A/Y395A mutations on ETS1Δ335 DNA binding stoichiometry. We find that, identical to wild-type ETS1Δ335, ETS1Δ335 R394A/Y395A completely protects both EBS core sequences in wild type MMP3 from digestion by DNAse I (Figure 4B). We also observe several enhancements of cleavage that are characteristic of ETS proteins binding to EBSs in footprinting assays (2,20,21). Regardless, the ETS1Δ335 R394A/Y395A -mediated protection of the 5’ EBS appears at substantially lower protein concentration than that at the 3’ EBS in comparison to the protections seen with ETS1Δ335 WT (compare Figure 2A to Figure 4B). Thus, the ability of ETS1Δ335 to cooperatively bind to two EBS DNA in a cooperative fashion is apparently disrupted by the R394A/Y395A mutations. However, these mutations do not block the ability of the protein to dimerize, as an ETS1Δ335 R378C/R394A/Y395A variant can still be crosslinked by BMOE both in the absence or presence of various DNA sequences (Figures 4C and figure 4D). Addition of DNA does not appear to alter the amount of dimer formed in comparison to that formed in the absence of DNA, however.

### Mutation in Helix H1 Blocks DNA Binding but not Dimer Formation

We showed previously that an L337A mutation blocks the formation of ETS1-p51 dimers, both with and without DNA (2). This residue is part of helix H1, which is thought to allow ETS1 to non-specifically probe the DNA phosphate backbone (22). As ETS1-p51 dimerizes only in the presence of DNA but ETS1Δ335 forms dimers in absence of DNA (2), we sought to determine if this mutation has similar effects on DNA binding and dimerization of ETS1Δ335.

ETS1Δ335 L337A is unable to form a complex with any DNA in an EMSA (Figure 5A). However, as noted previously (23), meaningful but unstable complexes are occasionally undetectable using an EMSA. Therefore, we also assayed the interaction of ETS1Δ335 L337A with the MMP3 sequence by DNase I footprinting. We find that the L337 mutation apparently blocks the interaction of ETS1Δ335 with DNA of any kind; ETS1Δ335 L337A neither protects the EBSs in MMP3 from cleavage nor are cleavage enhancements characteristic of ETS1-DNA complex formation observable (20,21)(Figure 5B).

**Figure 5:**
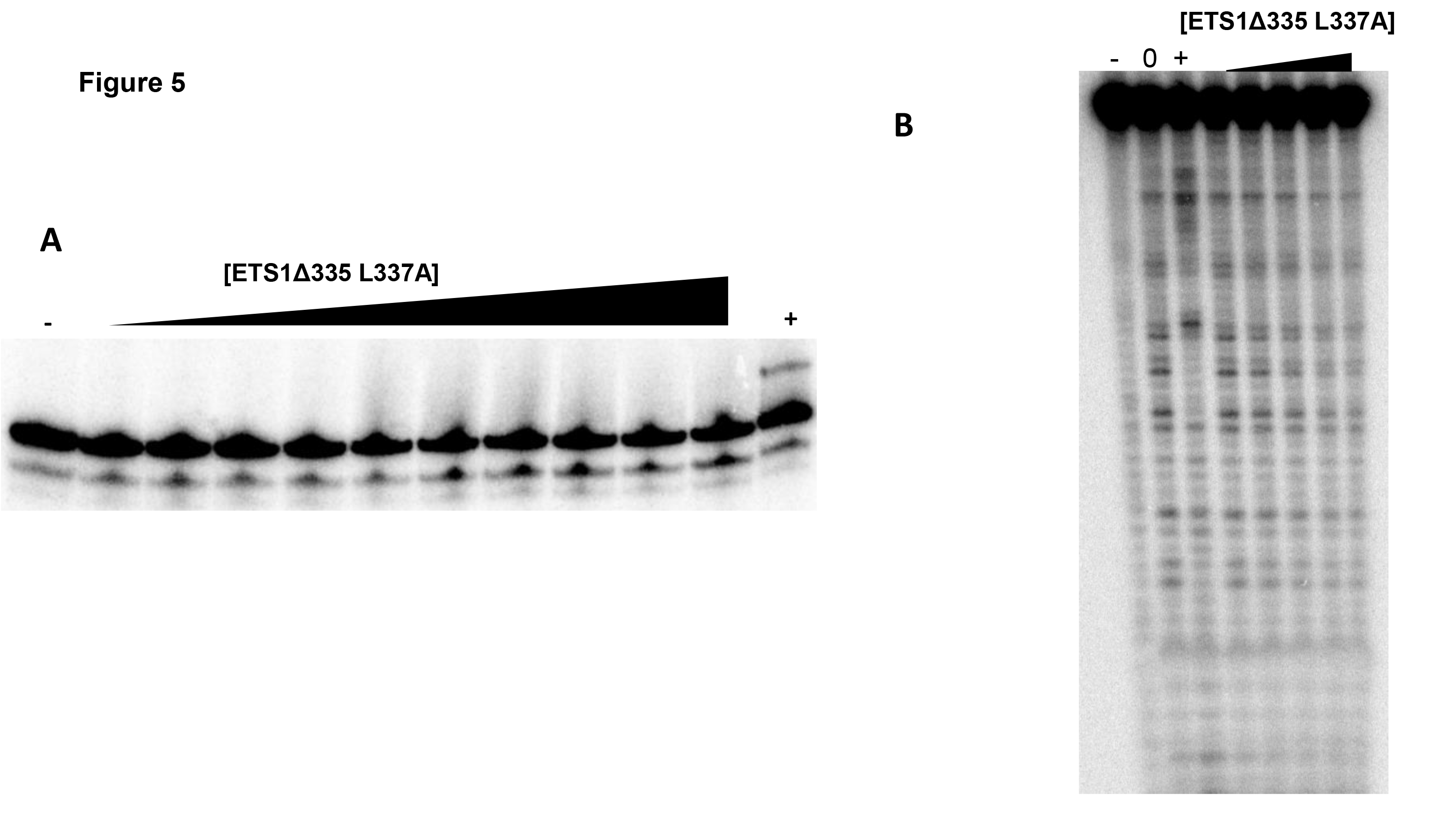
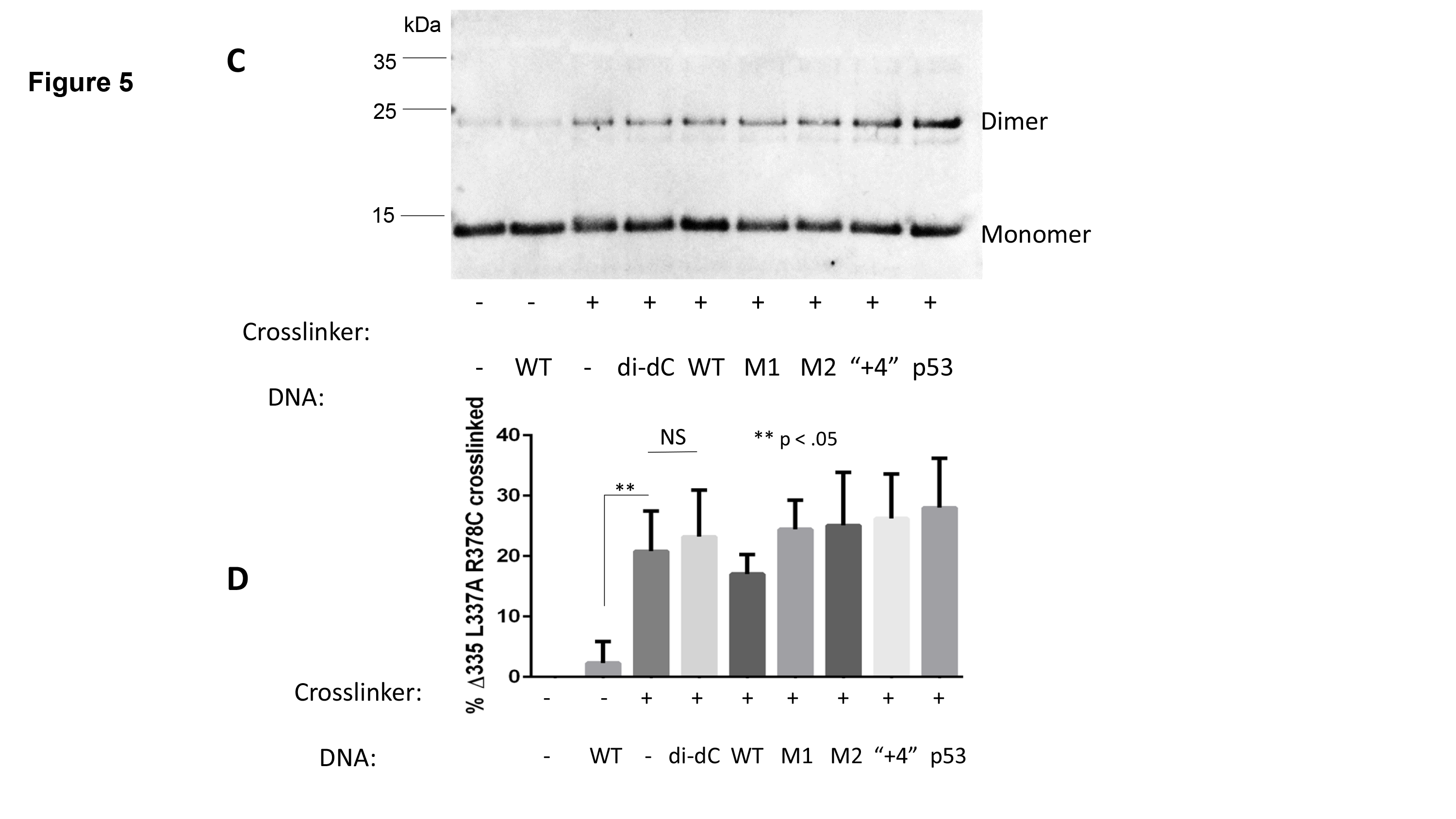
DNA Binding and dimer formation of ETS1Δ335 L337A. (A) Increasing concentrations of ETS1Δ335 L337A were mixed with radiolabeled binding site containing, MMP3 WT. Protein-DNA complexes were resolved via native PAGE as described in Methods and Materials. Shown is a Phosphorimager scan of the gel. Protein-DNA complexes are denoted with asterisks. “-”indicates no protein added. (B) Radiolabeled DNA bearing MMP3 wild type were incubated with increasing amounts of ETS1Δ335 L337A. Complexes were then subjected to DNase I cleavage and the resulting DNA fragments were resolved via denaturing PAGE. Shown is a Phosphorimager scan of the gel. The lane labeled witha ‘-’ contains uncleaved DNA and the lane labeled with 0 contains DNA incubated with DNAse I in the absence of protein. Positions of bases protected from cleavage by ETS1Δ335 L337A binding are bracketed. Enhancements of cleavage are noted with asterisks. (C) Shown is an immunoblot of ETS1Δ335 L337A crosslinked by BMOE in the presence or absence of the indicated DNA sequences. Positions of protein monomers and dimers are indicated. (D) Quantification of (C). Error bars represent standard deviation derived from four or more replicate experiments.

As the L337A mutation prevents ETS1-p51 DNA binding and dimerization, we sought to determine whether the L337A mutation had any effect on ETS1Δ335‘s ability to dimerize. To do this we examined the ability of ETS1Δ335 L337A R378C mutant protein to form crosslinkable dimers. When assayed by BMOE-mediated crosslinking, this protein also forms dimer complexes in the absence of DNA (Figure 5B and 5C). Adding different sequence DNAs had no significant effect on the amount of dimer formed by of ETS1Δ335 bearing the L337A mutation.

### Measurement of ETS1Δ335 Association State by FRET Analysis

Previous reports suggested that ETS1Δ335 stably binds DNA as a monomer (12), while work described here shows that ETS1Δ335 forms dimers both in the presence and absence of DNA (Figure 3). Therefore, we hypothesized that DNA acts to catalyze the separation of ETS1Δ335 dimers into monomers, which then stably associate with DNA. To test this idea, we used FRET to directly examine the effect of added DNA on the oligomeric state of ETS1Δ335. For these experiments, we labeled the N-terminal primary amine of ETS1Δ335 with either Alexa Fluor 488 or a fluorescent quencher, 4-((4-(dimethylamino)phenyl)azo)benzoic acid, succinimidyl ester, or dabcyl, which absorbs light but does not itself emit. In this assay, separation of dimers into monomers will allow an increase in fluorescence intensity, where formation of dimers will decrease it.

When a population of Alexa 488-labeled ETS1Δ335 is incubated with equimolar dabcyl-labeled ETS1Δ335, the fluorescence emitted at 525nm (Alexa 488’s emission peak) drops by approximately 25% in comparison to the same amount of Alexa 488-labeled ETS1Δ335 alone. This observation confirms that ETS1Δ335 dimerizes in the absence of DNA. As increasing amounts of MMP3 are added the fluorescence intensity increases, rising above that seen in the absence of DNA, before plateauing at saturating concentrations of DNA (Figure 6). Adding either MMP3 M2, which contains only one EBS, or p53 DNA, to which ETS1Δ335 does not bind, both induce an increase in fluorescence (Figure 6). This finding suggests that regardless of its sequence, DNA catalyzes the separation of ETS1Δ335 into monomers.

**Figure 6:**
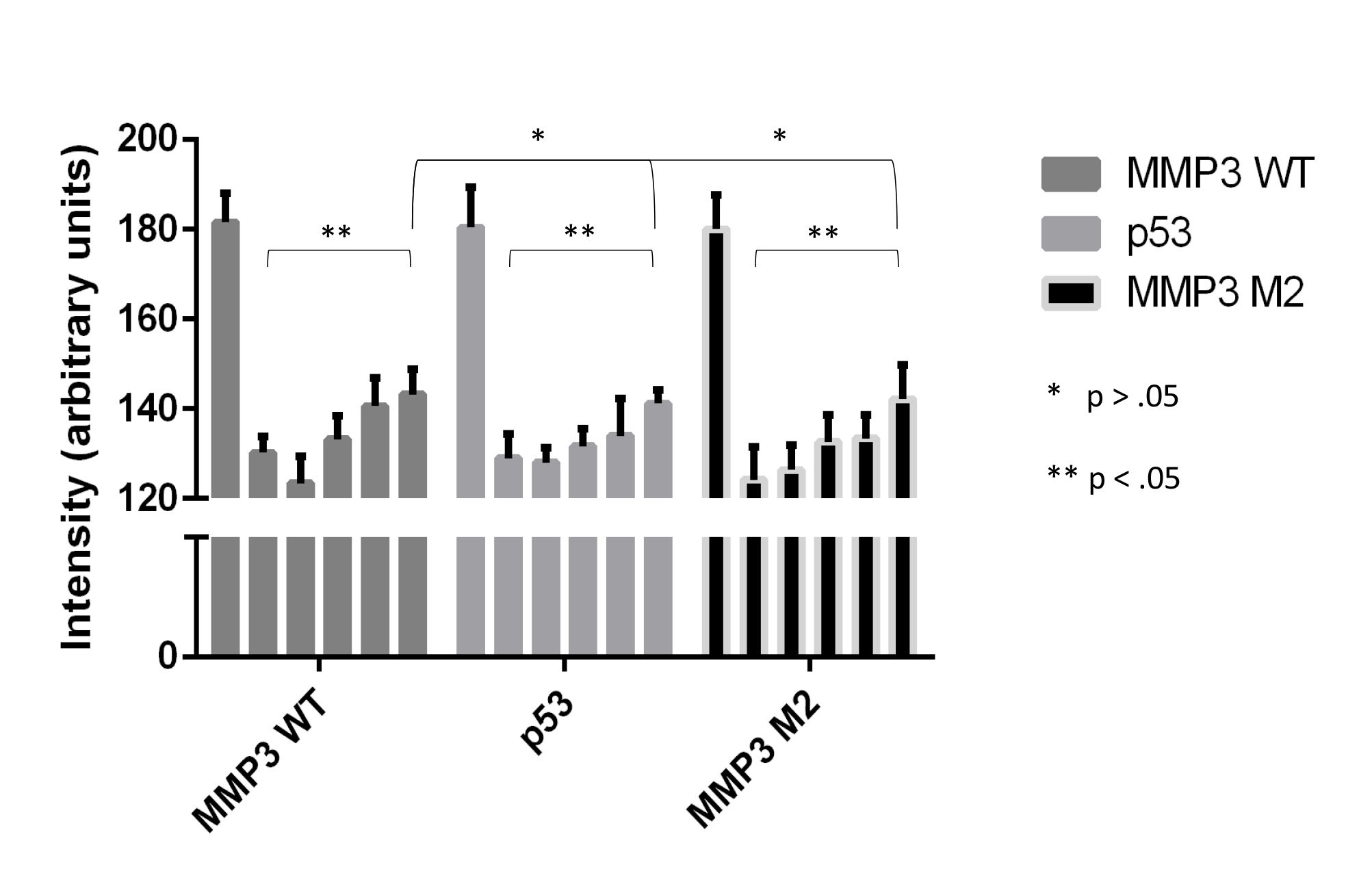
Fluorescence analysis of ETS1Δ335 association state. Two populations of purified of ETS1Δ335 were N-terminally modified with either Alexa Fluor 488 or a fluorescent quencher, dabcyl. The intensity at 525nm of Alexa-labeled ETS1Δ335 alone was assayed (first bar in each panel). Following this, equimolar dabcyl-labeled ETS1Δ335 was added and the intensity measured again (second bar). Increasing amounts of different DNA sequences were titrated in and the intensity assayed again, with the final titration being a saturating concentration of DNA. Error bars are standard deviation.

## Discussion

As the majority of ETS family proteins bind to extremely similar DNA sequences (24), one of the overarching goals in understanding ETS function is to determine how individual proteins in the family are each able to regulate a unique set of promoters (16). Two explanations have been suggested previously: indirect readout of DNA, where, non-conserved, non-contacted bases differentially affect an ETS transcription factor’s affinity for a binding site (2), and protein-protein interactions, where association with unique homo- or heterologous partner proteins causes structural rearrangements that modulate a protein’s ability to bind to DNA (13,16,25).

The questions of how ETS proteins exhibit differential functions extend to splice variants of a single ETS protein as well. When assayed *in vitro,* two of the three reported splice variants of ETS1, ETS1-p51 and ETS1-p42, display essentially identical DNA binding preferences (26,27). However, ETS1-p51 and ETS1-p42 have different gene regulatory activities *in vivo* (28). In light of this paradox, studies on ETS1-p42 have focused on the consequences of the loss of the autoinhibitory module on oligomer formation(19), without examining the biochemical nature of its DNA-binding properties. Due to the loss of the autoinhibitory module, it has been largely assumed that ETS1-p42 binds with 1-to-1 stoichiometry. This assertion is supported by static light-scattering data indicating that it binds to DNA as a monomer (12).

Nonetheless, several lines of evidence are inconsistent with the suggestion that ETS1-p42 binds as a monomer. We showed that ETS1Δ335, an N-terminal deletion that recapitulates ETS1-p42 binding, forms dimers in the absence of DNA (2). We also showed that autoinhibition in the full-length variant of ETS1 prevents self-association in the absence of DNA (2). Additionally, a study on the related ETS protein PU.1 suggests that, despite also seemingly binding DNA as a monomer, that protein binds to DNA in a negatively cooperative fashion, forming what the authors termed an “asymmetric dimer” (20). “DNA-free” dimerization is a property shared by other ETS proteins. Notably, the ETS TF ELK1 forms dimers when localized in the cytoplasm (29). This dimerization is thought to protect the protein from proteasomal degradation before it can cross over into the nucleus, where it preferentially accumulates. Once there, ELK1 apparently binds DNA as a monomer (29,30).

In this work we show that, despite apparently binding as a monomer to all binding sites tested in EMSAs, ETS1Δ335 also binds cooperatively to a DNA sequence containing two EBSs. That is, mutation of one of two EBSs in a DNA sequence substantially reduces the affinity of ETS1Δ335 for DNA. (Table 1).The suggestion that ETS1Δ335 binds cooperativity is supported by DNase I footprinting analysis which shows protection from cleavage at multiple EBS sites simultaneously (Figure 2A). Similarly, the protection persists even if one of the two EBSs present in the MMP3 sequence is mutated (Figures 2B and 2C). In addition, crosslinking analysis similar to that reported previously (2) shows that ETS1Δ335 forms dimers both in the absence and presence of DNA, with a statistically significant drop in dimer formation observed only in the presence of what is apparently a high affinity half-site (Figure 3).

The defects in cooperativity and DNA-binding seen in the R394A/Y395A and L337A mutants, respectively, still require explanation. Considering that the R394A/Y395A mutation disrupts the ability of ETS1-p51 to bind to DNA cooperatively, we suggest these mutations have similar effects on ETS1Δ335. In this case, cooperative binding to a high affinity site (the upstream EBS still present in MMP3 M2) which facilitates binding to a low affinity site (the downstream EBS still present in MMP3 M1) is disrupted. Binding to the high affinity site does not apparently require an intact helix H3, while the low affinity site does. When present together, both EBSs and helix H3 facilitate highest affinity binding to MMP3 wild-type site (compare Table 1 and Table 2).

As ETS1Δ335 R394A/Y395A mutant protein is still able to bind to DNA, albeit at a lower affinity and with a DNAse I footprinting pattern similar to that of wild-type ETS1Δ335, it appears that the R394A/Y395A mutation disrupts ETS1Δ335‘s ability to bind DNA cooperatively. This loss of cooperation between individual ETS1 monomers is reminiscent of the effect that the R394A/Y395A mutations have on ETS1-p51. In that protein, these mutations disrupt ETS1’s ability to cooperate and counteract autoinhibition, resulting in the formation of DNA-free dimers (2).

In ETS1-p51, an L337A mutation blocks the ability of the protein both to bind to DNA and to form dimers (2). However, when assayed by CD spectroscopy, both wild-type and ETS1-p51 L337A mutant display a similar loss of helical content upon addition of DNA (2). Therefore, we conjecture that ETS1Δ335 L337A separates dimers into monomers normally, but cannot meaningfully interact with DNA.

Consistent with this idea, the results of our FRET investigation indicate that in the absence of DNA, ETS1Δ335 is a dimer (Figure 6). Adding DNA of any sequence apparently catalyzes the dissociation of this protein from dimers into monomers (Figure 6). Thus, our FRET results appear to support a model where ETS1Δ335 binds to DNA as a monomer. The observed DNA binding cooperativity and dimer formation seen in the EMSAs, footprinting, and crosslinking assays support a model where ETS1Δ335 binds stably to DNA as a monomer, but also as an unstable dimer. This is particularly evident in footprinting assays that show two protections from cleavage when only one EBS is present (Figures 2B and 2C). This would be similar to the “asymmetric dimer” model proposed for the related ETS protein PU.1 (20). However we did not detect a decrease in fluorescence at the highest concentrations of DNA,in our FRET assay, a result which would be consistent with the formation of the “asymmetric dimer” proposed to occur in PU.1 (20). We suggest that the FRET assay, while sensitive to separation of dimers into monomers, is insensitive to formation of an unstable “asymmetric dimer”, possibly because the distance between to two fluors in the complex is too large to allow FRET.

Regardless, if, as seen in the FRET assay, DNA stimulates the separation of ETS1Δ335 dimers into monomers, (Figure 6), why does DNA have little to no effect on dimer formation when assayed by crosslinking (Figure 3)? We suggest that the results seen in Figure 3 represent a dynamic equilibrium between unbound dimers and bound monomers. As individual proteins shift from monomers to dimers they become crosslinked, removing them from the equilibrium. In the FRET assay, this does not occur, and in the presence of excess DNA, the equilibrium is biased towards the formation of ETS1Δ335 dimers.

Taken together, our data supports a new model for ETS1-p42 DNA-binding. In the absence of DNA, ETS1-p42 likely exists as an unbound dimer. Upon interaction with DNA, the dimers separate (Figure 6), with one protein binding stably as a monomer. As ETS1-p42 has been shown to bind DNA in heterodimers (11,25), it is likely that upon separation of unbound ETS1-p42 dimers, a heterodimer can form if a partner protein is present. However, if a heterologous partner protein is not available, a second ETS1-p42 monomer can associate with the DNA-bound ETS1-p42 monomer, forming a dimer that is structurally dissimilar to the unbound dimer. We suggest that the phenomenon of ETS1-p42 forming unbound homodimers is a form of autoinhibition that enables ETS1-p42 to form DNA-bound dimers upon separation of the unbound dimer. This alternate form of autoinhibition could have several potential functions. If it is similar to ETS1-p51 autoinhibition, it likely acts to prime ETS1-p42 for potential heteromeric complex formation. Such partnerships have been observed in with a number of different proteins (11,25,31). It may act as a form of protection against proteasomal degradation while traversing the cytoplasm, as seen in ELK1 (29). Additionally, it may act to facilitate the cooperative binding at single EBSs that we observe here.

As autoinhibition in full-length ETS1 appears to function to shield a protein-dimerization interface, so too does dimerization in this splice variant. This is evident as ETS1Δ335 dimers can be crosslinked via BMOE by the same residue, R378, as full-length ETS1. This residue exists in the cleft between two ETS1 proteins bound to a head-to-head binding site. This finding shows that both proteins form dimers using the same interface.

